# *In vitro* survival and neurogenic potential of central canal-derived neural stem cells depend on spinal cord injury type

**DOI:** 10.1101/2024.01.27.577563

**Authors:** Lars Erik Schiro, Ulrich Stefan Bauer, Christiana Bjorkli, Axel Sandvig, Ioanna Sandvig

## Abstract

The central canal (CC) of the spinal cord is a neurogenic niche consisting of quiescent neural stem cells (NSCs) capable of responding to traumatic damage to the spinal cord by increasing their proliferative activity and sending migrating progeny toward the site of injury, where they contribute to the formation of the glial scar. However, CC NSCs have been demonstrated to have the capability to differentiate into all neural lineage cells *in vitro*, but also *in vivo*, in response to infusion of specific growth factors that promote neuronal induction after injury, as well as when transplanted into other neurogenic niches, such as the subgranular zone of the hippocampus. This suggests that CC NSCs may represent a recruitable endogenous source of neural lineage cells that could be harnessed to replenish damaged or lost neural tissue after traumatic spinal cord injury (SCI).

NSCs isolated from the CC neurogenic niche of uninjured rats and mice have been shown to display limited proliferative capacity *in vitro*, with significantly greater proliferative activity achieved with NSCs isolated from SCI-lesioned rats and mice indicating an injury-specific activation of the quiescent CC NSC pool. A central question that currently remains unanswered is whether, and to what extent the CC niche can spontaneously generate viable neurons, and act as a potential source of new cells to replace lost neuronal populations *in situ*, and whether SCI sequalae impact future NSC neurogenic potential. To address this question, we need to understand whether the nature of the injury plays a role in the CC neurogenic niche response. In this study, we compared the intrinsic proliferative response and neurogenic potential of NSCs harvested from the CC neurogenic niche in adult female Sprague Dawley rats by culturing said NSCs across three conditions; (i) control, i.e., uninjured tissue, (ii) after *in vivo* compression injury 3 days before harvesting, and (iii) after *in vivo* simulated burst fracture injury 3 days before harvesting *in vitro*. We found that lacerations of the dura mater surrounding the spinal cord during a compression injury resulted in drastically altered and persistent *in vitro* NSC behavior encompassing both proliferation and development compared to uninjured control and compression injury with the dura intact.

## Introduction

Traumatic spinal cord injuries (SCIs) tend to cause permanent, debilitating deficits and affect approximately 250,000-500,000 people annually worldwide, with 40-80 cases per 1,000,000, and a 2:1 ratio of men affected compared to women (WHO, 2013). Traumatic injuries represent up to 90% of all SCIs and consist primarily of localized compression of the spinal cord, hyperextension, and cracked or shattered vertebrae lacerating the dura mater, i.e., burst fracture injuries (Mehdar et al., 2019; Nobunaga et al., 1999; Patek & Stewart, 2023; Singh et al., 2003; Wilcox et al., 2004). Most traumatic SCIs occur in adolescence (women), early adulthood (men), and old age (both genders), with a significant impact on mental health, societal cost, and general quality of life, increasing early mortality risk by 2-5 times (WHO, 2013). Full recovery is only possible in very mild cases of SCI, however, some recovery of function may occur even after severe injury, except for complete transection of the spinal cord (Khorasanizadeh et al., 2019).

Several potential therapeutic avenues have been proposed to address the issue of functional recovery following SCI, each achieving limited success in mammalian animal models. Such approaches include surgical decompression (Lee et al., 2018; Rahimi-Movaghar, 2005), pharmacotherapy (Gazdic et al., 2018), implanted functional electrical stimulation systems (Formento et al., 2018; Hardin et al., 2007; Wagner et al., 2018), exogenous stem cell transplantation (Assinck et al., 2017; Coutts & Keirstead, 2008; Gong et al., 2020), and activation and reprogramming of endogenous NSCs (Kameda et al., 2018; Li et al., 2016; Venkatesh et al., 2019). Currently, surgical decompression is the only treatment employed in the clinic, however, its degree of success is highly dependent on the severity of the spinal cord compression injury and the timing of intervention (Badhiwala et al., 2021; Batchelor et al., 2013; Shields et al., 2005).

The candidate endogenous NSC population for several of these proposed therapeutic interventions resides within the canonical spinal cord neurogenic niche, consisting of a thin layer of NSCs lining the central canal (CC) running along the entire rostral-caudal axis of the spinal cord (Fabbiani et al., 2020; Schiro et al., 2022; Taupin & Gage, 2002). This neurogenic niche is phylogenetically conserved across multiple species including mammals and amphibians, albeit with greater regenerative potential in response to traumatic injury in lower-order vertebrates compared to higher-order vertebrates, particularly humans (Hugnot & Franzen, 2011; McDonough & Martínez-Cerdeño, 2012). The spinal cord CC neurogenic niche remains the least studied of the canonical neurogenic niches present within the central nervous system, compared to the subventricular zone (SVZ) and subgranular zone (SGZ) neurogenic niches (Hugnot & Franzen, 2011; Taupin, 2006).

In adulthood, under homeostatic conditions, the NSCs surrounding the CC of the spinal cord are characterized by slow, continuous proliferation, exclusively generating new NSCs in an apparent quiescent self-renewal state (Johansson et al., 1999). However, in response to a traumatic spinal cord compression injury, the rate of NSC proliferation is reduced within the first 24 hours after injury (Takahashi et al., 2003), but subsequently increases, peaking at a 50-fold increase approximately 72 hours post-injury. This peak lasts for about a week before slowly returning to baseline levels over the following 3-4 months (McDonough & Martínez-Cerdeño, 2012). During this injury-induced spike in proliferative activity, NSC progeny migrate towards the injury site, where they continue to proliferate and subsequently differentiate into reactive astrocytes, contributing to the formation of the glial scar (Nicaise et al., 2022), sealing the injury site and limiting secondary injury effects to protect surrounding healthy tissue. The glial scar constitutes a key barrier to outgrowing axons from the injured pathways from reconnecting with their original targets, limiting functional recovery (Bradbury & Burnside, 2019; Clifford et al., 2023; Leal-Filho, 2011; McDonough & Martínez-Cerdeño, 2012; Meletis et al., 2008; Moreno-Manzano, 2020; Nicaise et al., 2022; Tran et al., 2022).

Microenvironmental injury sequelae to SCI with compression injuries which leave the dural sack intact, and burst fracture injuries, which induce dural laceration and subsequent intradural decompression, may not be identical. A difference in the NSC response to lesion type has previously been observed between compression and transection injuries (complete and incomplete, (McDonough & Martínez-Cerdeño, 2012; Meletis et al., 2008). Thus, the CC neurogenic niche may also respond differently to compression and burst fracture injuries in a similar, or altogether different manner. In this context, a better understanding of the role of the endogenous spinal cord neurogenic niche response to compression and burst fracture SCI is highly relevant, as it may provide useful insights towards harnessing the endogenous neurogenic niche potential to promote repair.

In this study, we applied our recently developed protocol (Schiro et al., 2022) to isolate and culture NSCs from the CC neurogenic niche of uninjured control, compression (cSCI), and burst fracture SCI (bfSCI) lesioned rats. Our primary focus was on NSC survival and proliferative potential during *in vitro* expansion, and spontaneous structural organization during differentiation and neural cell maturation. We found that these *in vitro* properties of CC-derived NSCs differ largely depending on injury type, resulting in significantly different survival and proliferative activity between the two injury conditions and the uninjured control NSCs, in addition to drastically affecting both survival and spontaneous self-organization when differentiated towards a neural lineage fate (neurons and astrocytes/glial cells).

## Materials and methods

### Animals

All animal procedures were approved by the Norwegian Animal Research Authority and in accordance with the Norwegian Animal Welfare Act §§ 1-28, the Norwegian Regulations of Animal Research §§ 1-26, and the European Convention for the Protection of Vertebrate Animals used for Experimental and Other Scientific Purposes (FOTS ID: 18066 & 24506). The animals were kept on a 12 h light/dark cycle under standard laboratory conditions (19–22°C, 50%–60% humidity), and had *ad libitum* access to food and water. All animals used were 10-week-old female Sprague Dawley rats (Janvier Labs, Le Genest-Saint-Isle, France), housed at the Comparative Medicine core facility, NTNU. A total of 39 animals were used in this study, divided into three groups: Group 1 – Uninjured Controls: n=16, Group 2 – Compression spinal cord injury (cSCI): n=12, Group 3 – Burst fracture spinal cord injury (bfSCI): n=11.

### Surgical induction of spinal cord injury

Each animal was anesthetized before the surgical procedure and anesthesia was maintained with 2%-3% isoflurane (Baxter, ESDG9623C) in 40% oxygen and 60% nitrogen gas mix, with body temperature regulated by a heating pad beneath the animal. Eye-protective gel (Viscotears, 7939) was applied to the animaĺs eyes immediately after being positioned for surgery. The animal’s back was shaven clean before a local injection of 0.1 ml Marcaine (2,5 mg/ml: Aspen Pharma Trading Limited, Ireland) was administered subcutaneously at the site of surgery as a local analgesic in addition to 0.45 ml Temgesic (0.03 mg/ml, Indivor Europe Limited) administered intraperitoneally as a general analgesic for intraoperative pain relief. After the thoracic level (T)10 vertebra was located by counting spinal processes along the rostral-caudal axis of the spine from the base of the skull (Watson et al., 2009), a 2 cm long incision was made from T8 to T12. The dorsal spinal column (T9-T11) was partially exposed, and the spinous process of T11 was removed for easier access. The ligamentum flavum was carefully transected at the caudal base of the T10 lamina, before complete removal of the dorsal T10 Lamina, exposing the spinal cord. For the cSCI, an arterial clip was gently placed onto the spinal cord, covering approximately ¾ of the dorsal/ventral axis before clamping to ensure direct lesioning of the central canal, while leaving some ventral spinal cord tissue intact. The arterial clip was kept in place for 60 seconds to induce the compression injury. For the bfSCI, a small cut was made to the dura mater to simulate laceration of the dura by intruding bone fragments. An arterial clip was then immediately placed onto the spinal cord covering the laceration to simulate the temporarily increased pressure from the injury. The arterial clip was removed after 60 seconds, and the surgical wound was sutured. After the lesioning procedure was completed, the animals were allowed to wake up in a heated cage before being transferred to a single-housing cage once consciousness was regained. Over the next 3 days post-surgery, the health of the animals was closely monitored, and the bladder was manually expressed every 6-8 hours. 0.45 ml Temgesic (0.03 mg/ml, Indivor Europe Limited) was administered intraperitoneally as a general analgesic to manage pain every 6-8 hours for the first 48 hours for all operated animals, and for longer if deemed necessary, based on the Rat Grimace Scale (Sotocina Susana & Sorge Robert, 2011).

### Tissue processing and immunohistochemistry

For labeling of proliferating cells in the CC neurogenic niche after injury, each animal was injected intraperitoneally with 37 mg/kg BW of 5-ethynyl-2′-deoxyuridine (EdU, ThermoFisher, A10044) at 48 hours after surgery, and kept alive for 24 hours before tissue fixation by intracardial perfusion. The animals were anesthetized with 4% isoflurane before an overdose of Pentobarbital (5 ml/kg Bodyweight, Ås Produksjonslab AS, 005608) administered by intraperitoneal injection. Each animal was then perfused by an initial 50 ml of heparin-supplemented saline (1:1000, LEO Pharma, 09837) solution at room temperature (23°C) at a rate of 20 ml/min, followed by 200 ml of chilled (4°C) 4% fresh paraformaldehyde (PFA, Sigma, P6148) at 20 ml/min. Once the perfusion was complete, the spinal cord was extracted and post-fixated overnight in 4% PFA at 4°C, before being transferred to a cryoprotective sinking solution (30% Sucrose (Sigma, 84097) in MQ water) and stored for 1 week at 4°C before sectioning. The spinal cords were sectioned coronally at 40 µm in 6 equally spaced series on a freezing microtome for immunolabelling and analysis.

One randomly selected series of tissue sections from each spinal cord were dehydrated in ethanol, cleared in xylene (VWR International, Radnor, PA, USA, 1330-20-7), and rehydrated before staining with Cresyl violet (Nissl; Sigma-Aldrich, St. Louis, MO, USA, C5042-10G) for 3 minutes on a shaker while protected from light. The sections were then alternatively dipped in ethanol–acetic acid (5 ml acetic acid in 1L 70 % ethanol) and rinsed with cold water until the desired differentiation was obtained, then dehydrated, cleared in xylene, and coverslipped with Entellan containing Xylene (VWR International, Radnor, PA, USA, 100503-868). One series of tissue sections from each spinal cord was left incubated in reaction buffer (Click-IT), CuSO4, Alexa Fluor Azide (488), and reaction buffer additive (Invitrogen, C10337) for 30 minutes for EdU labeling. The incubation was followed by a wash with 4′, 6-diamidino-2-fenylindol (DAPI; 1:10 000; Sigma-Aldrich, Saint-Louis, MO, USA, 28718-90-3) and phosphate buffer (PB) for 10 minutes, followed by a wash in PB for 10 minutes.

For immunohistochemistry (IHC), heat-induced antigen retrieval (HIAR) was carried out on two series of tissue (stained with Nestin/Ki67 and Iba1/GFAP) at 60 °C for 2 hours in (PB). Sections were subsequently washed 3 times for 10 minutes with PB containing 0.2 % Triton X-100 (Merck, Darmstadt, Germany, 108603; PBT). Next, sections were blocked using 5 % normal goat serum (Abcam, Cambridge, UK, ab7481) in PBT (PBT+) for 1 hour before incubation with the primary antibody. Sections were incubated with primary antibodies (see table S1 for primary antibodies used) in PBT+ for 4 hours at 4 °C. To label and visualize primary antibodies, directly conjugated Alexa Fluor secondary antibodies (Invitrogen, California, USA) at a 1:400 concentration were used (Alexa Fluor 488, 568, and 647, Table S1). First, sections were washed 3 times for 10 minutes with PBT. Then, sections were incubated with secondary antibodies for 2 hours at room temperature, protected from light. Then sections were washed for 10 minutes with DAPI (1:10 000) and PB, followed by 3 washes for 10 minutes with PB.

### NSC harvest and culturing

The procedures used for dissecting out the spinal cord, and harvesting and dissociating the NSCs for this study are described in detail in our published protocol (Schiro et al., 2022). See Figure 1 for the location of NSCs harvested from injured animals.

**Fig. 1:**
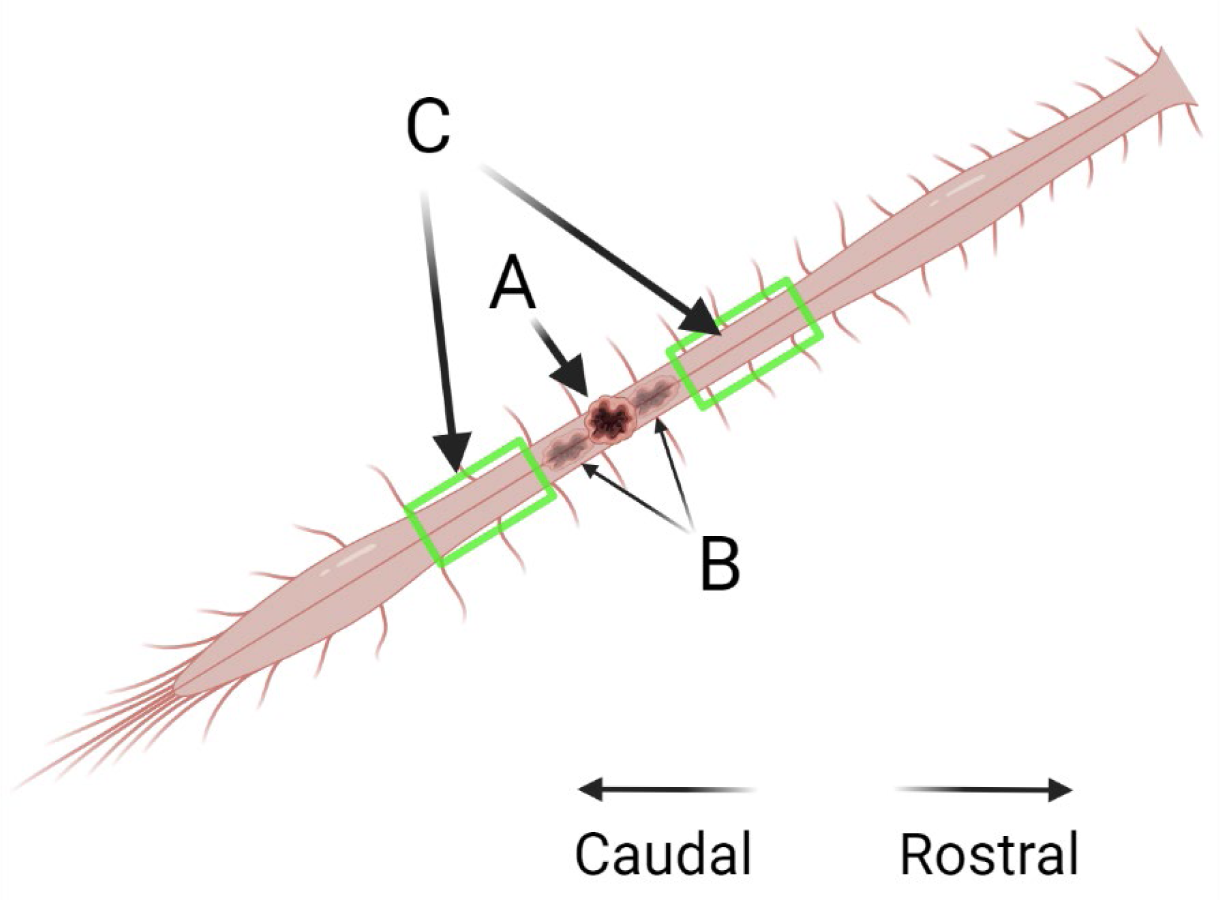
Schematic of NSC harvest location from injured animals. A: Primary injury site (T10). B: Extended damage of tissue from injury. C: Area excised for NSC harvest to avoid inclusion of injured/destroyed tissue into the cultures.

A total of 1×10^6^ live cells were seeded from each animal across 2x culture wells (5×10^5^/well) across the uninjured control cSCI and bfSCI conditions and expanded under near identical culturing conditions for 20 days (DIV), with culture media composition remaining the same for the full 20 days. The NSCs were passaged twice during expansion, first at 10 DIV (P1) and were split equally into 4 culture vessels (1x 6-well plate) to avoid a high degree of confluence (+90%) during P1, and second at 20 DIV (P2). All wells from each condition (plate) were pooled, dissociated, and counted together.

After 20 DIV, the NSC cultures were passaged and seeded onto Poly-L-Ornithine (Sigma-Aldrich, P4957) & Natural Mouse Laminin (Gibco, Cat# 23017-015) coated 13 mm^2^ coverslips (1×10^5^live cells/coverslip) and cultured for 3 days with identical media composition as during DIV 0-20.

At 23 DIV the NSC cultures were separated into two groups: Group 1: Fixed and immunolabelled for NSC culture characterization (see Table S1 for antibodies used). Group 2: Culture media changed to differentiation media (described in detail below).

Full culturing, expansion, dissociation, counting, and seeding procedures for the CC-derived NSCs are described in detail in our published protocol: Schiro, et al. 2022.

NSC cultures were differentiated for a total of 45 days, with 80% of media changed every 3 days and the cultures imaged every 15 days to track development (0, 15, 30, and 45). Differentiation base media: Neurobasal 1x base medium (Gibco, Cat# 21103049), 1x B27 Plus (Gibco, A3582801), 1x N2 (Gibco, Cat# 17502047), 1x L-Glutamine (Gibco, Cat# 25030081) and Heparin 2.5 µg/mL (Sigma-Aldrich, H3149). Differentiation supplements: BDNF (PeproTech, Cat# 450-02), CNTF (PeproTech, Cat# 450-13), GDNF (PeproTech, Cat# 450-51) and FBS (Sigma-Aldrich Cat# F9665). The differentiated cultures were fixed on day 45 (68 DIV) and immunolabelled for neural fate development (see supplementary table for antibodies used).

The methods used for EdU labeling, fixing, and immunolabelling the NSC and differentiated cultures for this study are described in detail in our published protocol; Schiro, et al. 2022.

### Imaging, quantification, and statistics

Live cells were imaged using a Zeiss Axiovert 25 inverted phase contrast microscope with a Zeiss Axiocam 105 color camera and transmitted white light as the light source.

Cell proliferation and survival: Total cell counts, and percentage of live cells were counted automatically following dissociation by staining the cells with Trypan blue and counting using a Countess™ II Automated Cell Counter (Invitrogen, AMQAX1000) and averaged for each animal after each passage.

Tissue sections were imaged using a Mirax-midi slide scanner and an Axiovert A1 fluorescent microscope (Zeiss, 503 mono camera) using either reflected fluorescence (fluorescence imaging) or transmitted white light (Cresyl Violet; Nissl) as the light source.

Immunolabelled cell cultures were imaged using an EVOS M5000 (Thermo Fisher) and counted using FIJI’s (ImageJ) automated cell counting function based on ICC images with a 1.0 mm^2^ field of view per image. Structural network nodes and connecting filaments were counted manually from random 1 mm^2^ sample images from each differentiated culture.

Statistical data was tested for normal distribution (Shapiro-Wilk) and equal variance (F-test (Snedecor & Cochran 1989)) before mean comparison using Student’s two-tailed unpaired t-test for normal distributed and small sample size data (n < 6 (De Winter 2019)) or Mann Whitney U for non-parametric data in STATA/MP 18.

## Results

### Histological verification of injury and neurogenic niche activation

Verification of the lesion was done by Nissl staining and immunohistochemical characterization of the inflammatory response to SCI by staining for astrocytes (GFAP) and microglia (Iba-1) in the uninjured control and injured spinal cords (Fig. 2 A&B).

**Fig. 2:**
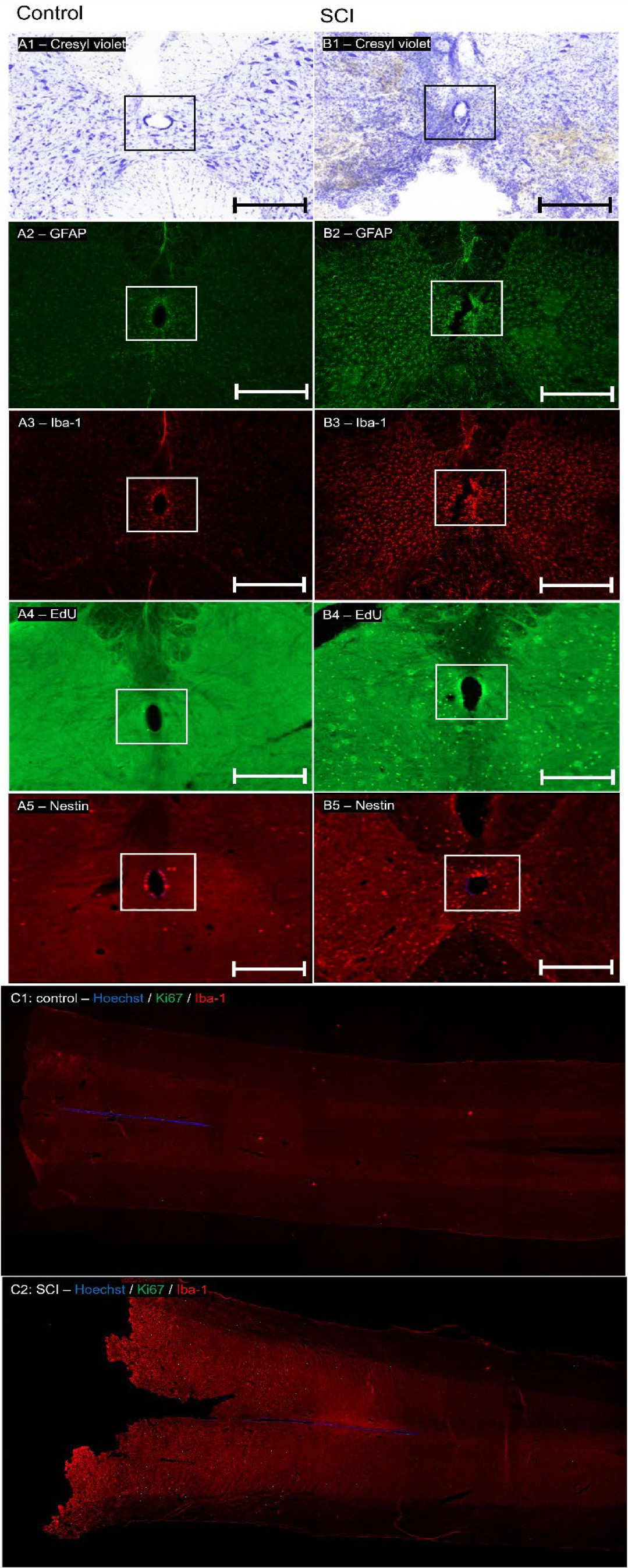
Confirmation of injury and neurogenic niche activation. A1: Histological Nissl-stained coronal section of uninjured control spinal cord tissue. B1: Histological Nissl-stained coronal section of injured spinal cord tissue. A2: Astrocytic (GFAP; green) expression in uninjured control spinal cord tissue. B2: Astrocytic (GFAP; green) expression in uninjured control spinal cord tissue in SCI lesioned spinal cord tissue. A3: Reactive microglia (Iba-1; red), expression in uninjured control spinal cord tissue. B3: Reactive microglia (Iba-1; red), expression in SCI lesioned spinal cord tissue. A4: Labelling of proliferative cell activity (EdU; green) in uninjured control spinal cord tissue. B4: Labelling of proliferative cell activity (EdU; green) in SCI lesioned spinal cord tissue. A5: Immunolabelling of Nestin-positive NSCs (red) in uninjured control spinal cord tissue (DAPI; magenta). B5: Immunolabelling of Nestin-positive NSCs (red) in SCI lesioned spinal cord tissue (DAPI; magenta). White/black box: Central canal, all scale bars: 200 µm. C1: Iba-1 (red) immunoreactivity and actively proliferating Ki67 positive (green) cells in the coronal section of the uninjured control spinal cord (DAPI blue). C2: Increased immunoreactivity of Iba-1 positive (red) reactive microglia and cell active Ki67 positive (green) proliferative activity in a coronal section of SCI lesioned spinal cord 3 days post-lesion (DAPI; blue). GFAP: Glial Fibrillary Acidic Protein, EdU: 5-ethynyl 2’-deoxyuridine, DAPI: 4’,6-diamidino-2-phenylindole.

Lesion-induced activation of the CC neurogenic niche resulted in a drastic, significant 378.5-fold increase in cell proliferation compared to the uninjured control, detected by positive 24h EdU labeling at 48-72 hours post lesioning (z_18_ = 3.833, p = 0.0001, Mann Whitney U, Fig. 2A&B4). In addition, a significant 34.9-fold increase in the number of Nestin immunoreactive cells was detected throughout the lesioned spinal cord compared to the uninjured controls (t_18_ = 5.5, p = 0.0003, unpaired two-tailed t-test, Fig. 2A&B5, Table S2).

### Initial NSC expansion as monolayer cultures

Microdissected NSCs from the CC of the spinal cord from all experimental conditions were initially cultured as free-floating cells after harvest (P0), which subsequently developed into free-floating neurospheres (Fig. 3A1, red arrows).

**Fig. 3:**
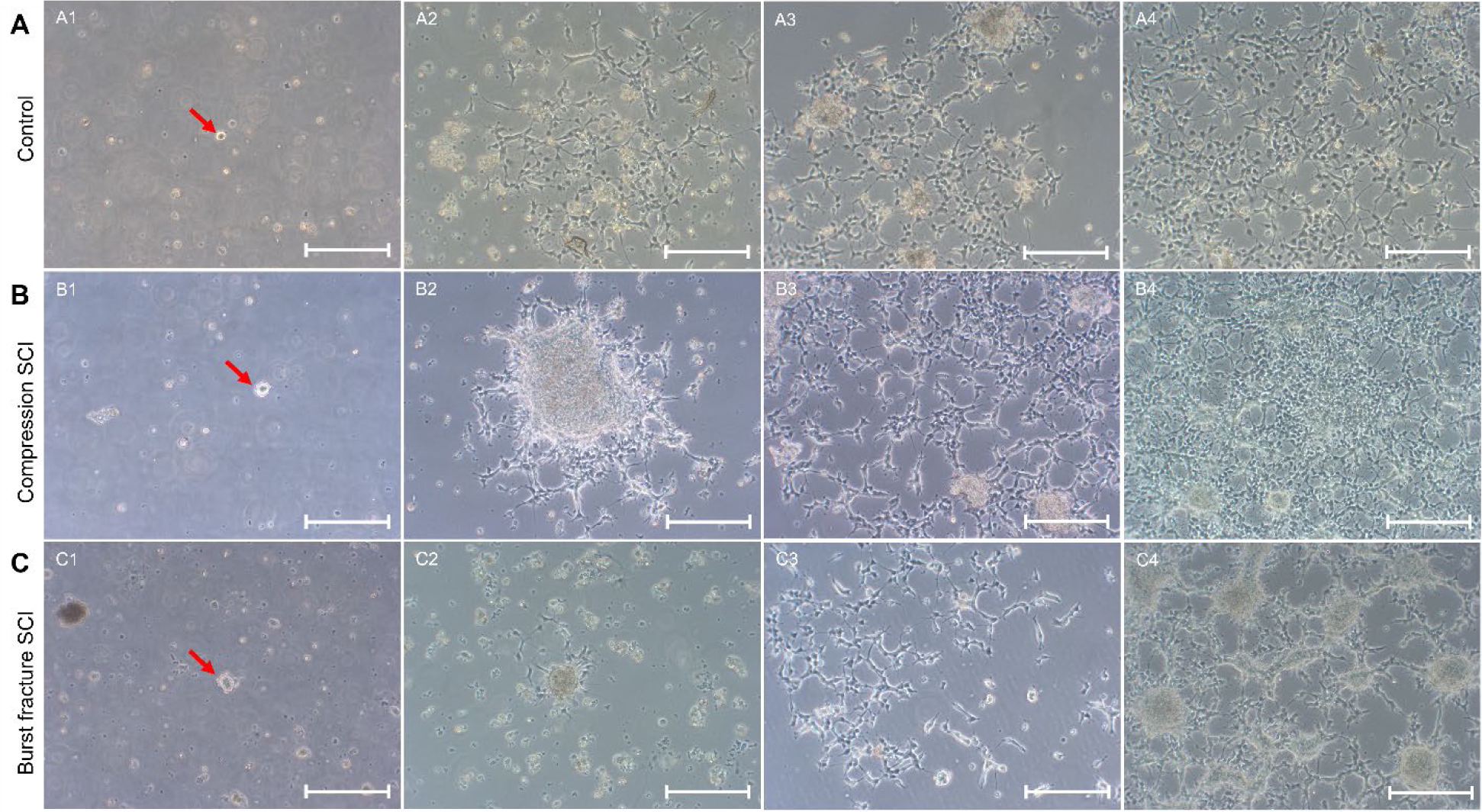
NSC *in vitro* profile over 20 days of *in vitro* expansion. Brightfield images of CC-derived uninjured control, cSCI, and bfSCI NSC culture development during expansion. A1: 4DIV free floating neurosphere (red arrow) of uninjured control NSCs. B1: 4DIV free floating neurosphere (red arrow) of cSCI NSCs. C1: 4DIV free floating neurosphere (red arrow) of bfSCI NSCs. A2: 10 DIV (pre-P1 passage) monolayer colony of uninjured control NSCs. B2: 10 DIV (pre-P1 passage) monolayer colony of cSCI NSCs. C2: 10 DIV (pre-P1 passage) monolayer colony of bfSCI NSCs. A3: 13 DIV monolayer colony of uninjured control NSCs. B3: 13 DIV monolayer colony of cSCI NSCs. C3: 13 DIV monolayer colony of bfSCI NSCs. A4: 20 DIV (pre-P2 passage) monolayer colony of uninjured control NSCs. B4: 20 DIV (pre-P2 passage) monolayer colony of cSCI NSCs. C4: 20 DIV (pre-P2 passage) monolayer colony of bfSCI NSCs. All scale bars: 200 µm. CC: Central Canal, cSCI: compression Spinal Cord Injury, bfSCI: burst fracture Spinal Cord Injury, DIV: days *in vitro*.

The free-floating NSC neurospheres from the uninjured controls spontaneously settled on the culture vessel surface after 5-7 DIV, developing into monolayer colonies (Fig. 3A2). This pattern repeated itself 3-5 days following the 1^st^ passage (13 DIV, Fig. 3A3). An initial drop in the number of live cells from harvest (P0) to P1 was observed, followed by a 12.2-fold increase in the number of live cells from P1 (10 DIV) to P2 (20 DIV, Fig. 3A4, 4A Table S3).

**Fig. 4:**
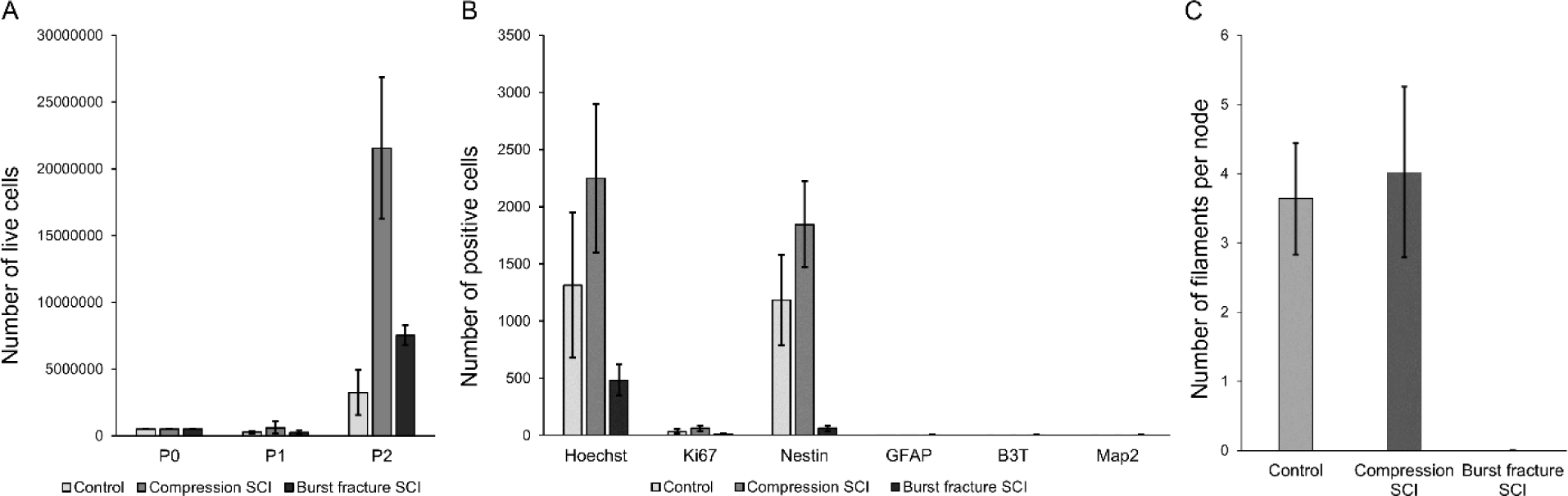
Proliferation and cell culture composition of CC-derived uninjured control, cSCI, and bfSCI NSCs following expansion. A: NSC cultures harvested from all 3 conditions (A) underwent limited proliferation during the first 10 days of expansion (P1). Proliferative activity was significantly greater across all 3 conditions during the following 10 days of expansion (10-20 DIV). B: Following expansion both the uninjured control and cSCI conditions consisted primarily of Nestin-positive NSCs, while the bfSCI cells were largely negative for the NSC marker. Active S-phase cell division took place within all 3 cultures (Ki67). Spontaneous neural differentiation was almost non-existent across all 3 conditions (glial: GFAP, neuronal: B3T, and Map2), however, a small but statistically significant higher level of spontaneous differentiation towards both glial (GFAP) and neuronal (B3T and Map2) cells was observed in the bfSCI condition compared to both the uninjured control and cSCI conditions. C: Number of cell body nodes interconnected by large glial and neuronal process filaments forming structural networks in the control, cSCI, and bfSCI NSC cultures after 45 dDiff. SCI: Spinal Cord Injury, NSC: Neural Stem Cell, B3T: Beta-III-Tubulin, Map2: Microtubule-Associated Protein 2, dDiff: days of Differentiation.

In contrast, cSCI NSCs typically remained clustered together after the free-floating neurospheres (Fig. 3B1, red arrow) spontaneously settled on the culture vessel surface on 5-7 DIV (P0), forming large, dense colonies surrounded by a small outer monolayer (Fig. 3B2). However, after the P1 passage (10 DIV) these cells formed more uniform monolayers after resettling (13 DIV, P1), similar to those of the uninjured control cultures, with few dense clusters throughout the culture (Fig. 3B3). The cSCI condition NSC cultures showed a small increase in live cells from harvest (P0) to P1, but not significantly different from the uninjured control condition (t_11_ = 1.93, p = 0.096, unpaired two-tailed t-test). This was then followed by a 35.4-fold increase in the number of live cells from P1 (10 DIV) to P2 (20 DIV, Fig. 3B4, 4A), a significantly greater increase than what was observed in the uninjured control cultures (t_8_ = 6.7, p = 0.004, unpaired two-tailed t-test, Table S3).

BfSCI NSCs formed few, dense colonies surrounded by individual cells spread out sporadically between the colonies once the free-floating neurospheres (Fig. 3C1, red arrow) spontaneously settled on the culture vessel surface on 5-7 DIV (P0, Fig. 3C2). Following the P1 passage (10 DIV), these NSCs resettled into a sparse monolayer on the culture vessel surface at 13 DIV (Fig. 3C3), before finally forming several dense colonies, with single NSCs interspersed between the larger colony clusters (Fig. 3C4). The bfSCI-derived NSCs showed a similar decrease in the number of live cells from harvest (P0) to P1 as the uninjured control cultures (t_8_ = 0.2, p = 0.84, unpaired two-tailed t-test), and lower than the cSCI condition (t_9_ = 1.9, p = 0.09, unpaired two-tailed t-test). This decrease was then followed by a 30-fold increase from P1 (10 DIV) to P2 (20 DIV), which significantly differed from both the uninjured control (t_8_ = 3.4, p = 0.008, unpaired two-tailed t-test) and cSCI condition cultures (t_6_ = 8.0, p = 0.0002, unpaired two-tailed t-test, Fig. 4A, Table S3).

### NSC immunoreactivity and spontaneous differentiation at 23 DIV

At 23 DIV (3 days post P2 passage (20 DIV)), NSCs from all 3 conditions were seeded at a density of 1×10^5^ cells per coverslip (13 mm^2^). Immunolabeling of the uninjured control condition cultures showed an average of 1,314 cells per mm^2^ as indicated by Hoechst positive nuclei, with 2.5% of the total cell population undergoing S-phase cell division at the time of fixation, as indicated by Ki67 immunoreactive nuclei. Nestin-positive cells made up 88.3% of the total cell population at 20 DIV (Fig. 4B, 5A1-2, Table S4). A significant increase (58.4%, z_58_ = 5.012, p < 0.0001, Mann Whitney U) in the total number of cells was observed in the cSCI cultures compared to the uninjured control, with a small but statistically significant greater number of cells (2.8%) undergoing S-phase cell division at the time of fixation as indicated by Ki67 positive nuclei (z_28_ = 3.762, p = 0.0002, Mann Whitney U). Nestin-positive cells made up 82.1% of the total cSCI cell population, a comparable proportion of the total cell population to what was found in the uninjured control condition (z_23_ = 0.684, p = 0.493, Mann Whitney U, Fig. 4B, 5A1-2, Table S4).

**Fig. 5:**
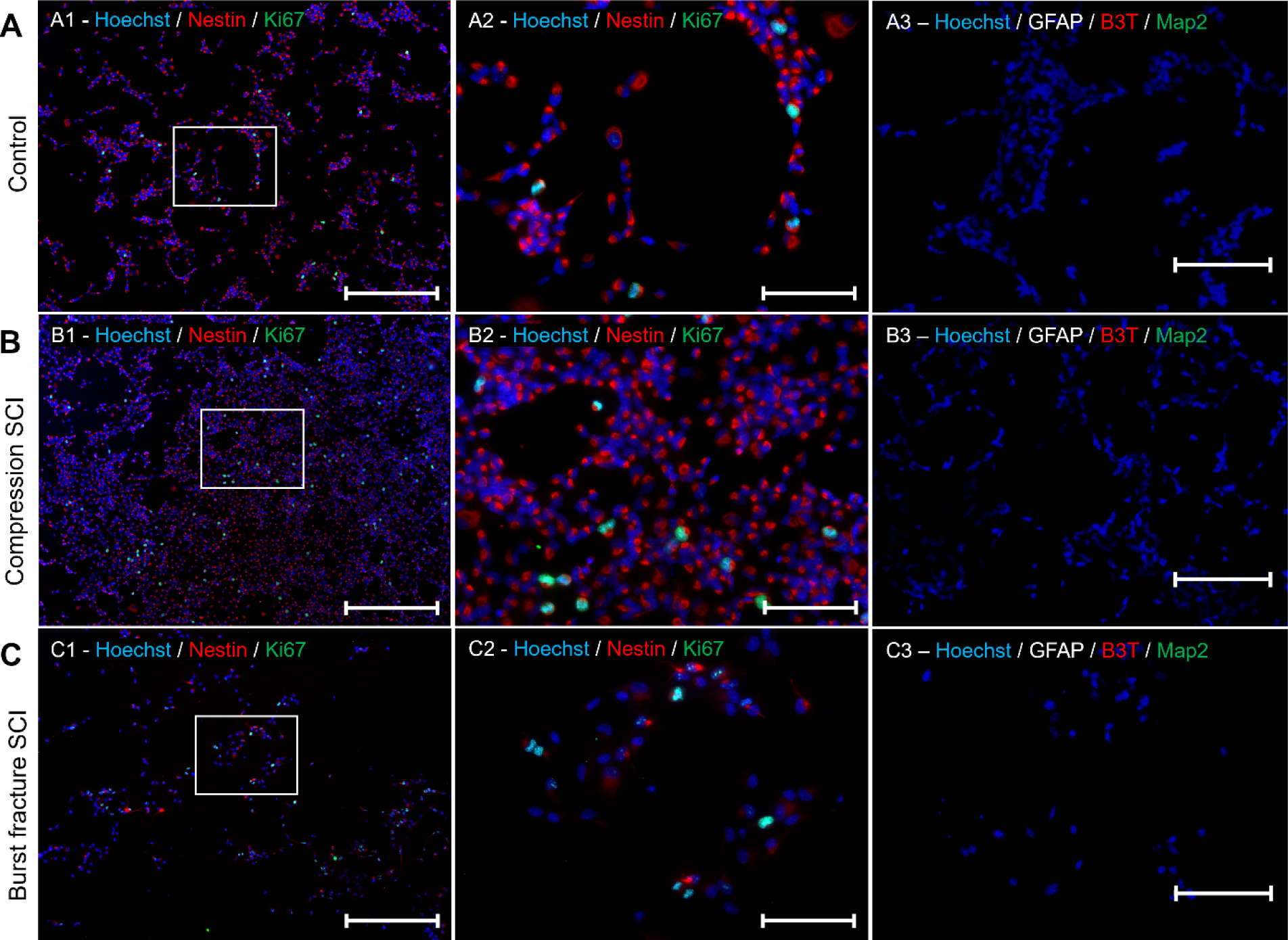
Expression of NSC- and proliferation markers after expansion. A1: Immunolabelling of actively proliferating (Ki67) NSCs (Nestin) in uninjured control cultures at 23 DIV. A2: Immunolabelling of actively proliferating (Ki67) NSCs (Nestin) in uninjured control cultures at 23 DIV (zoom white box A1). B1: Immunolabelling of actively proliferating (Ki67) NSCs (Nestin) in cSCI cultures at 23 DIV. B2: Immunolabelling of actively proliferating (Ki67) NSCs (Nestin) in cSCI cultures at 23 DIV (zoom white box B1). C1: Immunolabelling of actively proliferating (Ki67) NSCs (Nestin) in bfSCI cultures at 23 DIV. C2: Immunolabelling of actively proliferating (Ki67) NSCs (Nestin) in bfSCI cultures at 23 DIV (zoom white box C1). Blue: Hoechst, Red: Nestin, Green: Ki67, Scale bar 1: 200 µm, scale bar 2: 50 µm. A3: Immunolabelling of neural (GFAP, Beta-III-Tubulin and Map2) differentiation in uninjured control NSCs at 23 DIV. B3: Immunolabelling of neural (GFAP, Beta-III-Tubulin and Map2) differentiation in cSCI NSCs at 23 DIV. C3: Immunolabelling of neural (GFAP, Beta-III-Tubulin and Map2) differentiation in bfSCI NSCs at 23 DIV. Blue: Hoechst, Red: Beta-III-Tubulin, Gray: GFAP, Green: Map2, scale bar 3: 50 µm.

A significant decrease in the total number of cells was observed in the bfSCI condition compared to the uninjured control (63.2%, z_48_ = 5.565, p < 0.0001, Mann Whitney U) and cSCI (78.5%, z_48_ = 5.941, p < 0.0001, Mann Whitney U) conditions. Only 1.9% of all bfSCI cells underwent S-phase cell division at the time of fixation, a level comparable to both the uninjured control (t_23_ = 0.9, p = 0.36, unpaired two-tailed t-test) and cSCI (z_23_ = 1.833, p = 0.068, Mann Whitney U) conditions, as indicated by Ki67 positive cell nuclei. Only 16.7% of the total cell population in the bfSCI cultures were immunoreactive for Nestin, significantly lower compared to both the uninjured control (z_23_ = 4.160, p < 0.0001, Mann Whitney U) and cSCI (z23 = 4.161, p < 0.0001, Mann Whitney U) conditions (Fig. 4B, 5C1-2, Table S4).

Neither uninjured control nor cSCI condition cultures contained cells immunoreactive for neural lineage cells (GFAP, Beta-III-Tubulin, and Map2, Fig. 5A3-B3). However, a small portion of the bfSCI-derived NSC cultures underwent spontaneous differentiation with 0.4% GFAP, 0.6% Beta-III-Tubulin and 0.6% Map2 immunoreactive cells at 23 DIV (Fig. 4B, 5C3, Table S4).

### Self-organization of differentiated NSCs *in vitro*

Following expansion, the NSCs from all three conditions were cultured in expansion media for 3 days on PLO/laminin-coated glass coverslips before switching the cell culture media over to differentiation media. The NSCs of all 3 conditions settled onto the coverslips within these 3 days before spreading out into uniform monolayers after 3 days of differentiation (dDiff, 26DIV, Fig. 6A1-C1). At 15 dDiff the uninjured control NSCs had started developing into different, morphologically distinct cell types (Fig. 6A2) indicative of neural differentiation (S5). The cSCI NSC cultures displayed similar differential morphological development, in addition to a higher degree of clustering, compared to the uninjured control NSCs (Fig. 6B2). In the bfSCI NSC cultures however, we did not observe the same changes in cell morphology as in the uninjured control and cSCI NSC cultures, but instead an increase in debris from dead cells (Fig. 6C2). By 30 dDiff we observed a much greater degree of clustering in NSCs derived from both the uninjured control animals and after cSCI compared to 15 dDiff, with the cells now starting to self-organize into larger structures (Fig. 6A-B3). The bfSCI NSC cultures displayed minimal change on 30 dDiff compared to 15 dDiff, consistently appearing as large monolayer cultures with no signs of structural organization and minimal differentiation (Fig. 6C3).

**Fig. 6:**
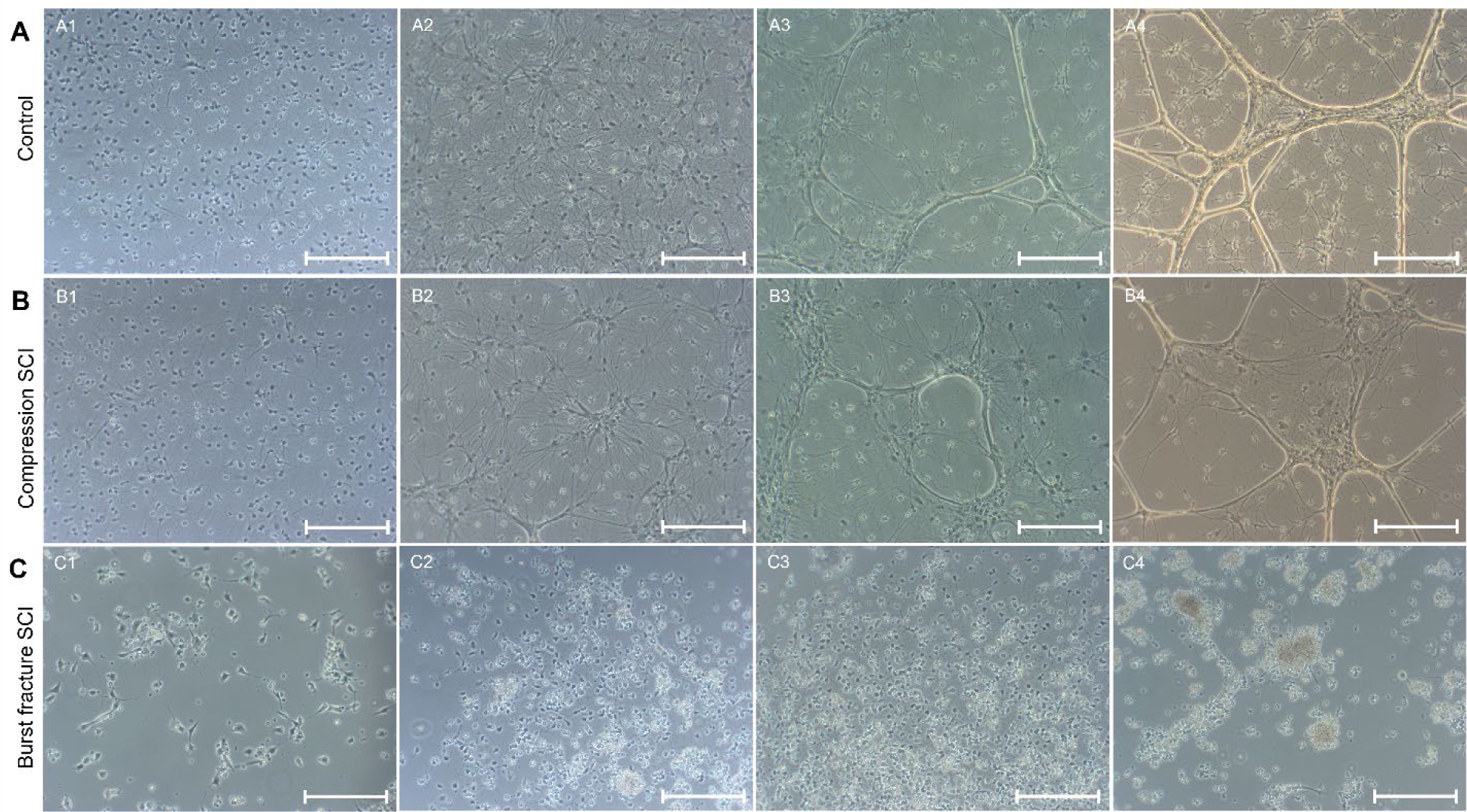
Self-organization of control and injury-activated NSCs *in vitro*. Brightfield images of CC-derived NSC cultures. A1: Uninjured control NSC culture after 3 dDiff. B1: cSCI culture after 3 dDiff. C1: bfSCI SCI NSC culture after 3 dDiff. A2: Uninjured control NSC culture after 15 dDiff. B2: cSCI culture after 15 dDiff. C2: bfSCI NSC culture after 15 dDiff. A3: Uninjured control NSC culture after 30 dDiff. B3: cSCI culture after 30 dDiff. C3: bfSCI NSC culture after 30 dDiff. A4: Uninjured control NSC culture after 45 dDiff. B4: cSCI culture after 45 dDiff. C4: bfSCI NSC culture after 45 dDiff. Scale bar: 50 µm. dDiff: days of differentiation.

At 45 dDiff, the uninjured control NSC cultures had organized into clearly defined structural networks (Fig. 6A4) consisting of nodes, mainly containing glial and neuronal cell bodies (Fig. 7A1-2, white box), interconnected by narrow filament-like GFAP positive glial structures containing Beta-III-Tubulin and Map2 positive neural cell processes (Fig. 7A1-2, red arrows). Each node was connected to an average of 3.63 other nodes (SD: 0.8, Fig. 4C) through these filaments.

The cSCI-derived NSC cultures followed a similar pattern of self-organization, with each node connected to an average of 4.02 other nodes (SD: 1.2, Fig. 4C), not significantly different from the uninjured control cultures (t_130_ = 1.93, p = 0.056, unpaired two-tailed t-test). However, only about 65% of the cSCI cultures fully organized into node-filament structural networks, with the remaining part of the culture consisting of widespread and connected GFAP positive glial cells and Beta-III-Tubulin and Map2 positive neuronal cells spread throughout (Fig. 6B4, 7B1-2).

The bfSCI-derived NSC cultures, however, did not organize into nodes and filaments like the uninjured controls and cSCI cultures but instead developed into smaller scattered colonies consisting of only a few glial GFAP positive and neuronal Beta-III-Tubulin and Map2 positive cells scattered throughout the culture vessel surrounded by debris (Fig. 4C, 6C4, 6C1-2), drastically differing in self-organization from the uninjured control and cSCI conditions with no nodes or connecting filaments formed.

## Discussion

Research on spinal cord transection injury in rodents has shown that the CC neurogenic niche response to injury, including NSC proliferation and migration capacity, is contingent on lesion severity. In spinal cord transection model injuries, the activation of NSCs appears localized, surrounding the site of the lesion (McDonough & Martínez-Cerdeño, 2012; McTigue et al., 2001; Meletis et al., 2008). Compression model injuries, such as the models used in this study, have been shown to elicit NSC activation along the entire rostral-caudal axis of the spinal cord. A key difference between transection and compression injuries is the rapid increase in intradural pressure from the lesioning process itself, and the subsequent swelling of the spinal cord resulting from hemorrhage and edema (McDonough & Martínez-Cerdeño, 2012; McTigue et al., 2001; Rowland et al., 2008; Tator & Fehlings, 1991; Zai & Wrathall, 2005). This leads to additional secondary injuries such as ischemia, inflammation, demyelination, and a greater extent of neuronal cell loss due to apoptosis and excitotoxicity compared to transection-type injuries (Hilton et al., 2017; Venkatesh et al., 2019). This increase in intradural pressure is absent in surgically transected spinal cords as the lesioning procedure requires opening the dural sack, cutting through the spinal cord as opposed to the application of external pressure (Cheriyan et al., 2014; Lukovic et al., 2015). As bone fragments can pierce and lacerate the dural sack in addition to compression of the spinal cord during burst fracture injuries, the increased pressure within the dural sack may be partially attenuated due to leakage of cerebrospinal fluid and neural tissue through the damaged dura during the initial compression event in this type of injury (Cammisa et al., 1989; Denis & Burkus, 1991; Park et al., 2011). This attenuation of intradural pressure during the traumatic event may potentially result in less extensive damage along the rostral-caudal axis of the spinal cord. However, intradural decompression may come at the cost of increased local tissue inflammation as blood and tissue may enter the dural sack through the lacerations (Kwiecien et al., 2020; Song et al., 2014), and cause greater severity of lesion cavitation and scar tissue invasion (Iannotti et al., 2006). This additional inflammatory reaction within the spinal cord may directly impact the NSC microenvironment and subsequent behavior such as observed in this study.

The NSCs used in this study were harvested at 3 days post-injury, a time point corresponding to the peak of proliferative activity within the CC proximal to the injury site after compression (Fig. 1 (Namiki & Tator, 1999)). We confirmed an increase in Nestin-positive cells throughout the gray and white matter of the spinal cord proximal to the injury site, compared to uninjured controls (Fig. 2A-B4-5). It has previously been reported that ablation of the Nestin-positive cells before lesioning resulted in significant worsening of pathological features such as demyelination, reduction of Olig2^+^ progenitors, and loss of neurons adjacent to the injury site in mice, indicating that Nestin-positive cells play an important role in the endogenous response to SCI (Cusimano et al., 2018). We observed an increase in proliferating cells as indicated by an increase of EdU-positive cells within the same region of the spinal cord (Fig. 2A-B4) following injury.

When culturing NSCs derived from the CC neurogenic niche of uninjured control, cSCI, and bfSCI, we observed significantly higher proliferative activity in the NSCs derived from cSCI animals compared to uninjured controls, as observed in previous studies (McDonough & Martínez-Cerdeño, 2012; Meletis et al., 2008; Namiki & Tator, 1999; Sabelström et al., 2014; Shu et al., 2022), but surprisingly, the proliferative activity of the cSCI NSCs was also significantly higher than that of the bfSCI derived NSCs. The proliferative activity of the bfSCI-derived NSCs was significantly higher than the uninjured controls, indicating an injury-related activation of the bfSCI NSCs. Comparison of the bfSCI NSCs with the cSCI derived NSCs however, indicated a significant differential activation of the neurogenic niche within the first 3 days post-injury, dependent on lesion type occurring, as also suggested by the subsequent behavior of the NSCs *in vitro* (Fig: 3 A).

It is not unlikely that ependymal cells constitute a large portion of the CC NSCs, however, we observed that uninjured control-derived NSCs remained positive for Nestin and SOX2 (Fig. 5B, 5A-C; Schiro et al. 2022), but were not positive for the ependymal marker FoxJ-1 (S7) (Li et al., 2018; Ren et al., 2017) or GFAP (Fig. 5) (McDonough & Martínez-Cerdeño, 2012) after 23 days of expansion. This indicates that the CC neurogenic niche derived NSCs are likely not ependymocytes or of ependymal descent and retain their stemness and proliferative potential (Nestin, Ki67, and EdU positive, Fig. 5, 7, Table S4.

**Figure 7:**
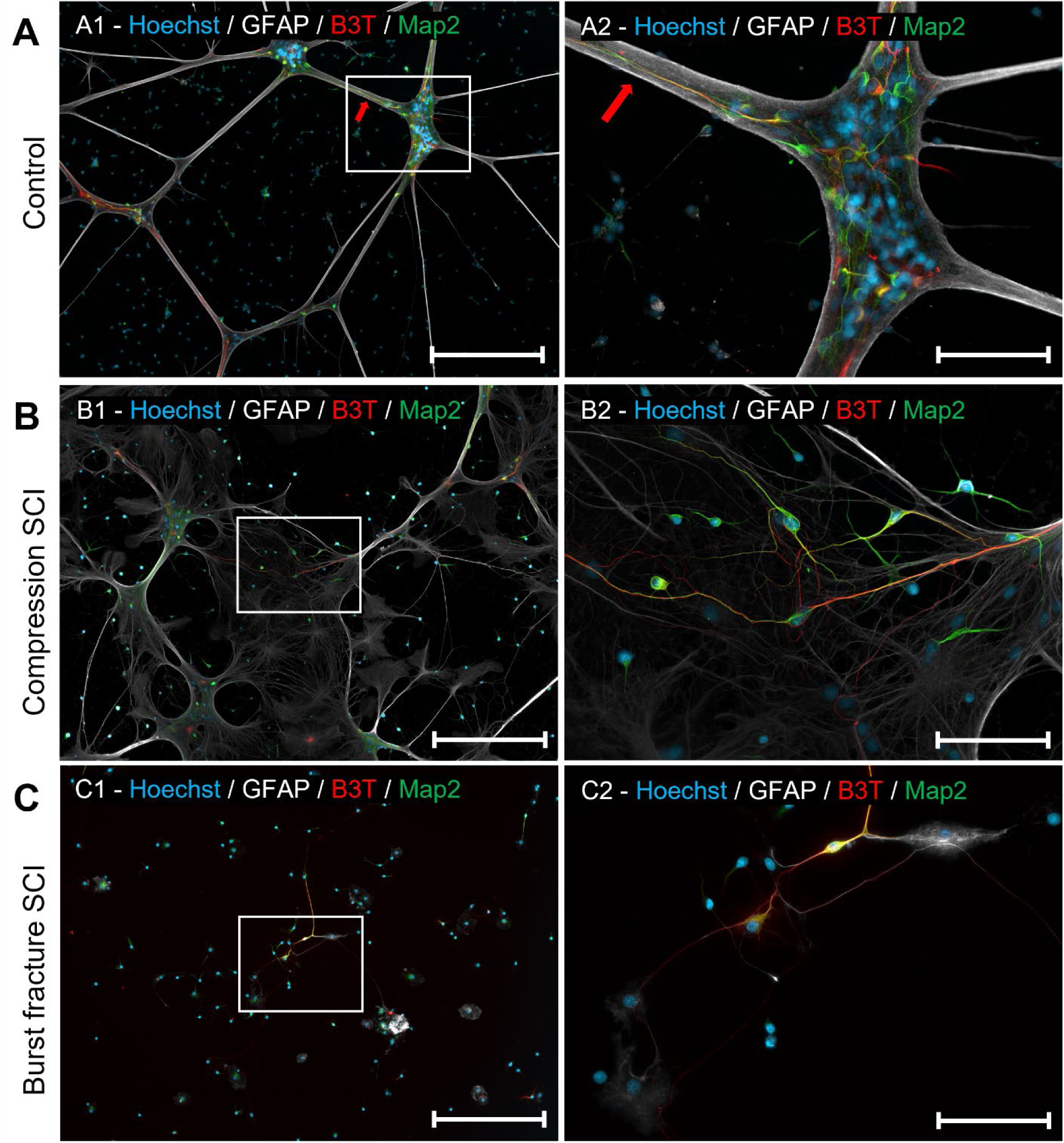
Immunocytochemistry of differentiated uninjured control and injury-activated central canal-derived NSCs. A: Self-organization of neuronal lineage cells derived from uninjured control NSCs into structural networks consisting of nodes (white box zoom: A2) interconnected by filaments (red arrows) after 45 days of differentiation. B: Self-organization of neuronal lineage cells derived from cSCI NSCs into semi-structured networks after 45 days of differentiation (white box zoom: B2). C: Small sporadic colonies of neuronal lineage cells differentiated from bfSCI NSCs after 45 days of differentiation (white box zoom: C2). Blue: Hoechst, White: GFAP, Green: MAP2, Red: Beta-III-Tubulin. Scale bar 1: 200 µm, scale bar 2: 50 µm.

As the CC-derived NSCs were able to differentiate into GFAP positive glial cells and Beta-III-Tubulin and Map2 positive neuronal cells (Fig. 7) *in vitro*, they likely represent a heterogenous NSC population that either no longer contain ependymal cells after 23 days in culture (Barnabé-Heider et al., 2010; Panayiotou & Malas, 2013), or FoxJ-1 is not required for maintenance of stem cell potential, as previously reported within the spinal cord neurogenic niche (Li et al., 2018).

Interestingly, while there was no significant difference in the number of Nestin-positive cells in the uninjured control and cSCI cultures compared to the total number of Hoechst-positive cells, the bfSCI NSC cultures contained a significantly smaller proportion of Nestin-positive cells compared to the total number of cells (indicated by Hoechst positive nuclear staining) and tended to cluster more together during expansion, as opposed to being spread out in a uniform monolayer like the uninjured control and cSCI NSC cultures (Fig. 3A-C1-4, 5A-C1-2). None of the NSC populations derived from any of the three conditions spontaneously differentiated towards a neural fate (Fig. 4B, 5A-C3). This suggests that the bfSCI NSCs may differentiate towards a non-neural fate *in vitro* as opposed to the uninjured control and cSCI NSC cultures, indicating a differential NSC activation from the altered microenvironment resulting from the dural laceration of the burst fracture SCI.

During neural differentiation, the *in vitro* behavior of the NSCs suggested a significant impact on SCI injury type. Specifically, the uninjured control NSCs self-organized into clearly defined nodes, primarily containing cell bodies, surrounded and interconnected by large GFAP-positive filament-like structures enwrapping dendritic and axonal neuronal processes (Fig. 6-7A), with observed instances of organization into larger structures (S8 A 1-3, red arrows) through the folding of large GFAP positive outer sheets over Beta-III-Tubulin and Map2 positive neuronal cells resting on top, forming large three-dimensional structures (S8 B 1-3, red arrows). The cSCI-derived NSCs developed along a similar path, partially self-organizing into large nodal structural networks similar to those observed in the uninjured control cultures. Large parts of the cSCI cultures also developed into large, unstructured GFAP-positive sheets with Map2 and Beta-III-Tubulin-positive neuronal cells sporadically spread throughout (Fig. 7B1-2). This difference in self-organization compared to the uninjured control-derived cultures was consistent across all experiments, suggesting a direct influence of the injury-specific *in vivo* signaling cues on the CC neurogenic niche and subsequently on the self-organizing behavior of these cells (Fig. 6-7B). The impact of the SCI on the development of the bfSCI NSCs however was much more prominent compared to the cSCI NSC cultures. Only a fraction of the bfSCI NSCs survived the differentiation process, resulting in too few remaining cells to organize into any form of structure. The result was sporadic GFAP positive glial and Beta-III-Tubulin and Map2 positive neuronal cells spread sparsely across the culture vessel and being detached from each other (Fig. 7C). Interestingly, many Hoechst positive cell nuclei were observed in the bfSCI cultures with no neural lineage immunoreactivity for GFAP, Beta-III-Tubulin or Map2 (Fig. 7C), suggesting that the altered NSC microenvironment resulting from dural laceration in a bfSCI may program the central canal NSCs to differentiate towards non-neural fate lineages.

## Conclusion

The injury type-dependent impact on NSC behavior persisted throughout expansion and differentiation under near identical culturing conditions and timings *in vitro* for up to 68 DIV and without any further external environmental cues save for those introduced to all conditions through the culturing cell media. This suggests that there is an early, robust differential activation of the NSCs residing within the CC neurogenic niche in response to different types of traumatic SCI.

This sensitivity of the CC neurogenic niche to injury and subsequent differential NSC response to a traumatic event may potentially play a role in why therapies targeting endogenous NSCs, such as pharmacotherapy, exogenous stem cell transplantation, and activation and reprogramming of endogenous NSCs, tend to fail to promote functional recovery following SCI. It is therefore also possible that with sufficient understanding of what drives and regulates the response of the CC neurogenic niche to different forms of SCI its resident NSCs may be induced to serve a direct and beneficial role in SCI rehabilitation.

## Author contributions

LES conceived the study, designed, and performed the experiments, collected, and analyzed the data, wrote, and finalized the manuscript; USB performed experiments, collected data, read, and edited the 17manuscript, CB performed the immunohistochemistry, Cresyl violet and relevant imaging, read, and edited the manuscript; AS provided funding and guidance, revised, and edited the manuscript; IS provided funding and guidance, revised, and edited the manuscript.

## Acknowledgments

The authors would like to acknowledge support by the Liaison Committee for Education, Research, and Innovation in Central Norway (Samarbeidsorganet HMN-NTNU), the Joint Research Committee between St. Olav’s Hospital and the Faculty of Medicine and Health Sciences (Felles Forskningsutvalg), NTNU, and Enabling Technologies, NTNU. We thank Marit Trones Rem for technical assistance.

## Supplementary

**Table S1:** Antibodies used for immunolabelling.

| Primary Antibody | Marker | Host | Concentration | Identifier |
| --- | --- | --- | --- | --- |
| Nestin | Neural Stem Cell | Mouse | 1:500 | ab6142 |
| Iba-1 | Microglia | Mouse | 1:500 | ab15690 |
| GFAP | Astrocytes | Rabbit | 1:1 000 | ab7260 |
| Ki67 | S-phase proliferating cell | Rabbit | 1:500 | ab15580 |
| EdU | Cells proliferating during exposure | - | - | C10337 |
| Map2 | Neuron | Chicken | 1:20 000 | ab5392 |
| Beta-2-Tubulin | Neuron | Mouse | 1:1 000 | ab78078 |
| FoxJ-1 | Motile cilia – Ependymal cell | Mouse | 1:100 | SC-53139 |
| GAP-43 | Growth cones | Rabbit | 1:250 | ab75810 |
| **Secondary Antibody** |  |  |  |  |
| 488 |  | Rabbit | 1:500 | ab150077 |
| 568 |  | Mouse | 1:500 | ab175473 |
| 647 |  | Chicken | 1:500 | ab150171 |
| Nuclear stain |  |  |  |  |
| Hoechst 33342 |  |  | 1:2 000 | C10337 G |
| DAPI |  |  | 1:10 000 | D1306 |

**Table S2:**
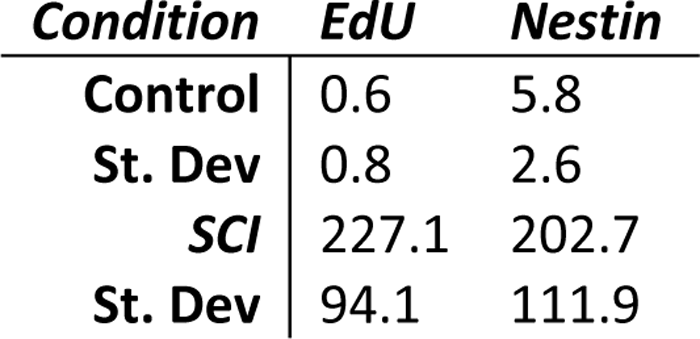
NSC *in vivo* proliferation data.

| Condition | EdU | Nestin |
| --- | --- | --- |
| Control | 0.6 | 5.8 |
| St. Dev | 0.8 | 2.6 |
| SCI | 227.1 | 202.7 |
| St. Dev | 94.1 | 111.9 |

**Table S3:** NSC expansion over 20 DIV.

| Niche | Generation | Average number of live cells | St. Dev |
| --- | --- | --- | --- |
| Control | P0 | 500 000 | 0 |
|  | P1 | 264 833 | 88 490 |
|  | P2 | 3 241 667 | 1 696 319 |
| cSCI | P0 | 500 000 | 0 |
|  | P1 | 607 714 | 457 828 |
|  | P2 | 21 550 000 | 5 282 992 |
| bfSCI | P0 | 500 000 | 0 |
|  | P1 | 250 675 | 137 299 |
|  | P2 | 7 535 000 | 728 320 |

**Table S4:** Average number of cells immunoreactive to proliferative, NSC, and neural fate markers.

| Condition | Hoechst | Ki67 | Nestin | GFAP | B3T | Map2 |
| --- | --- | --- | --- | --- | --- | --- |
| Control | 1 313.8 | 32.5 | 1 183.3 | 0.3 | 0 | 0 |
| St. Dev | 634.8 | 19.0 | 397.1 | 0.7 | 0 | 0 |
| cSCI | 2 247.6 | 62.7 | 1 844.6 | 0.1 | 0 | 0 |
| St. Dev | 648.6 | 22.6 | 376.0 | 0.3 | 0 | 0 |
| bfSCI | 475.7 | 9.4 | 63.6 | 1.8 | 3.1 | 2.7 |
| St. Dev | 137.6 | 6.3 | 25.0 | 2.0 | 1.4 | 1.9 |

**S5: Day 15 neural development of uninjured control culture.** Uninjured control culture neuronal cells developing after 15 days of differentiation. Blue: Hoechst, White: GFAP, Green: MAP2, Red: Beta-III-Tubulin. Scale bar: 50 μm.

**Figure S5:**
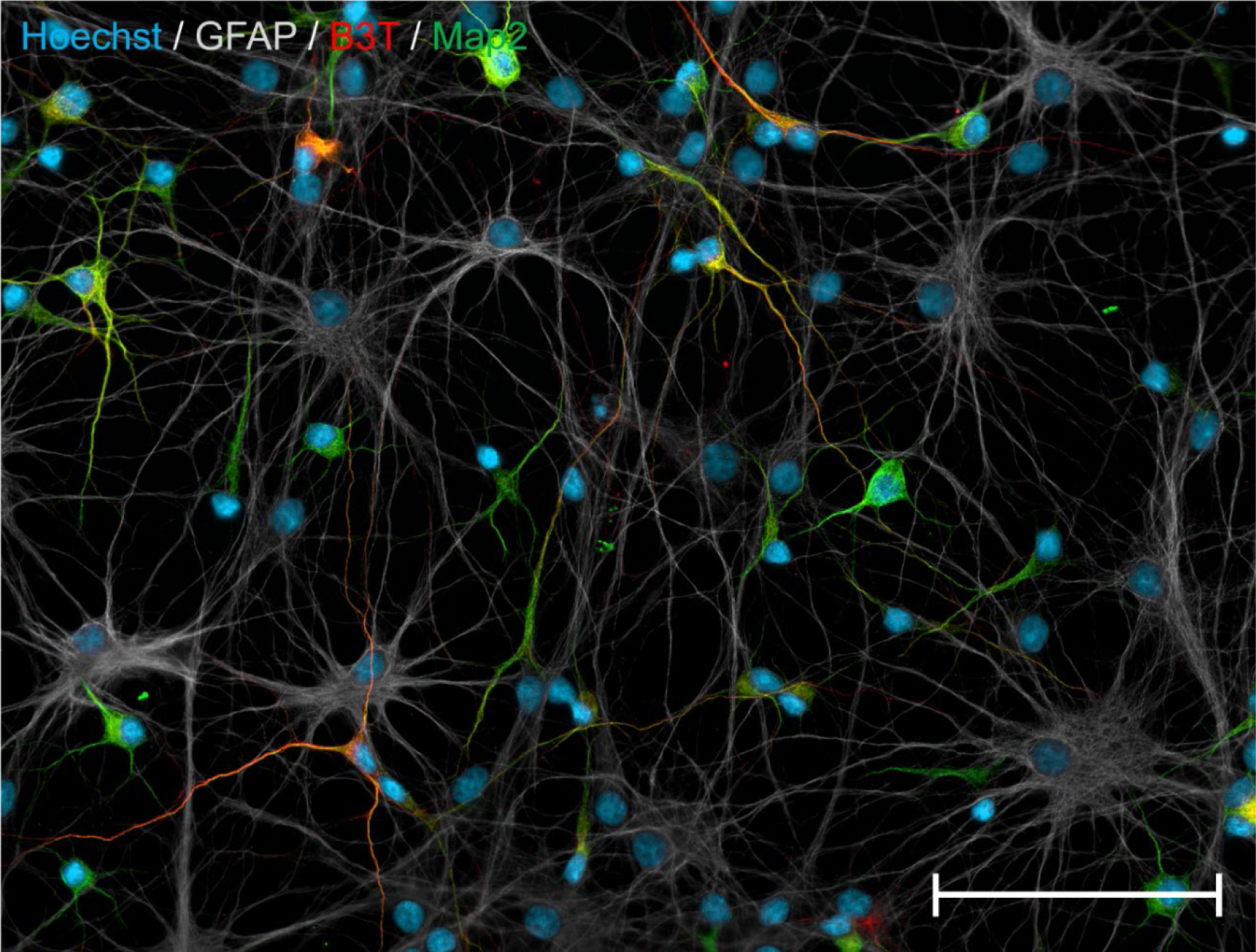
Day 15 neural development of uninjured control culture. Uninjured control culture neuronal cells developing after 15 days of differentiation. Blue: Hoechst, White: GFAP, Green: MAP2, Red: Beta-III-Tubulin. Scale bar: 50 µm.

**Figure S6:**
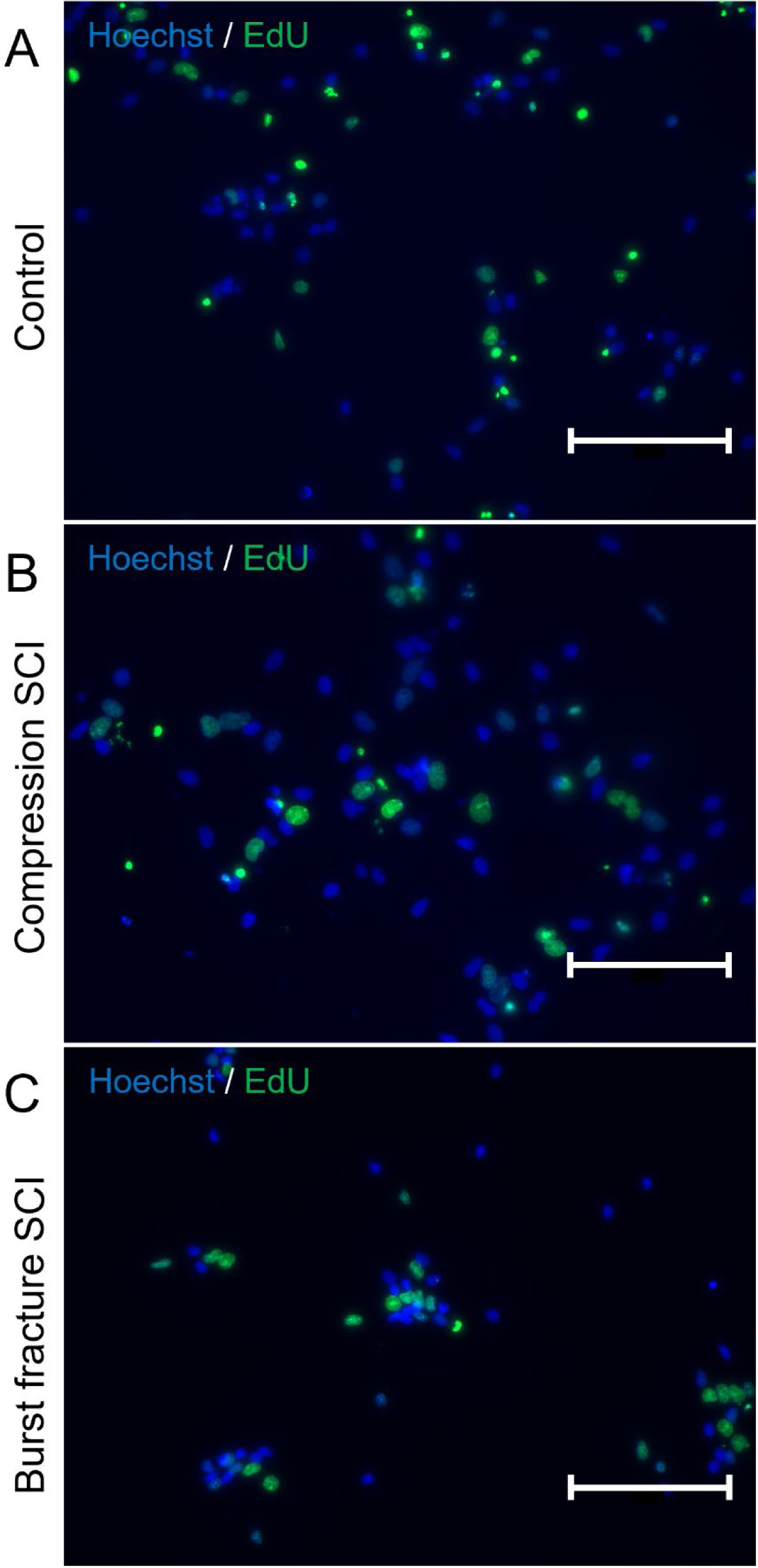
Confirmation of proliferation of central canal-derived NSC control and injury conditions after 23 days of expansion: A: 24-hour EdU labeling of proliferating NSCs in the uninjured control condition. B: 24-hour EdU labeling of proliferating NSCs in the compression SCI condition. C: 24-hour EdU labeling of proliferating NSCs in the burst fracture SCI condition. Scale bar: 250 µm.

**Figure S7:**
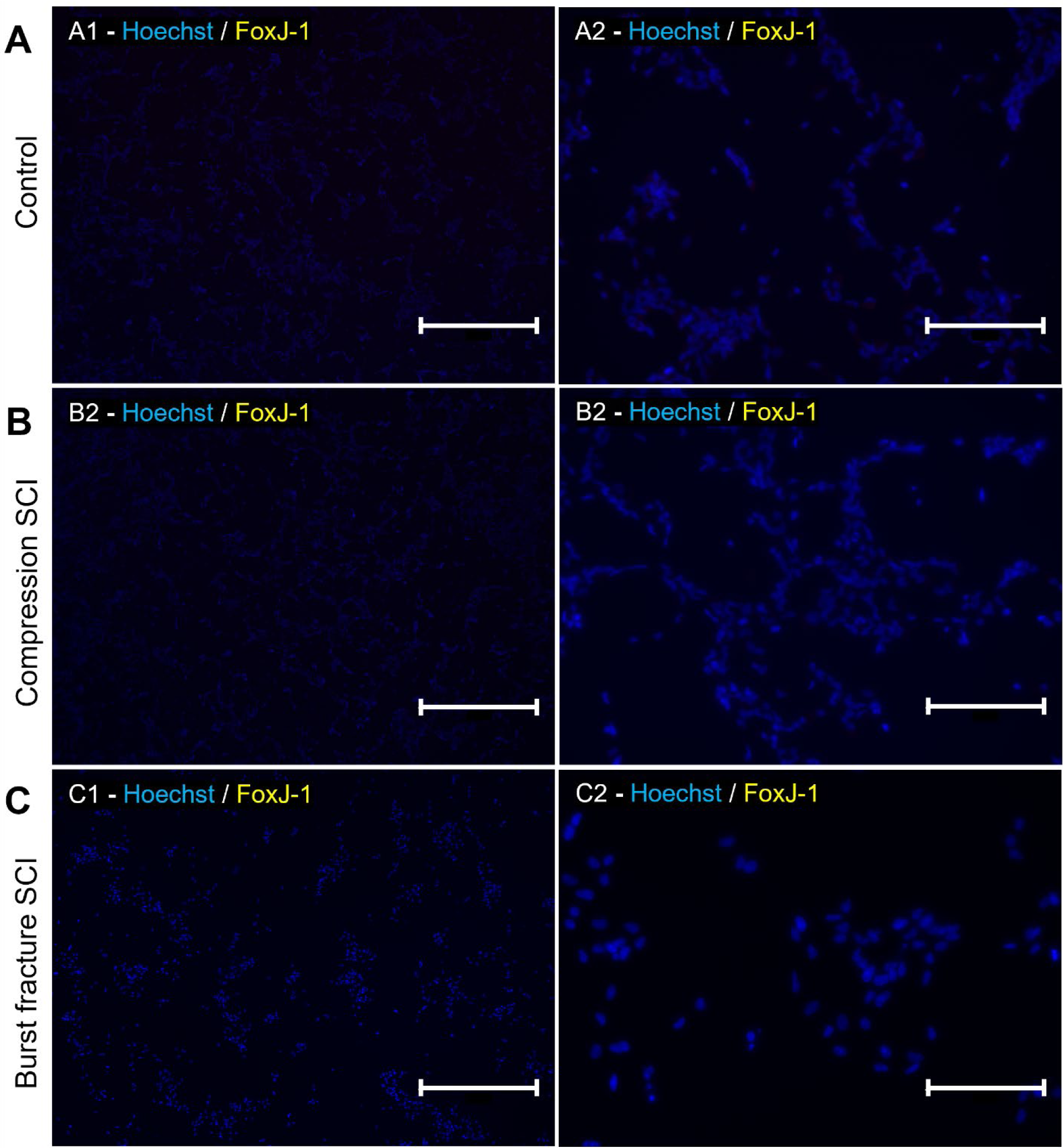
FoxJ-1 expression in central canal-derived NSCs after 23 days of expansion. A: Uninjured control NSC expression of FoxJ-1. B: Compression SCI NSC expression of FoxJ-1. Burst fracture SCI NSC expression of FoxJ-1. Scalebar 1: 1250 µm. 2: 250 µm.

**Figure S8:**
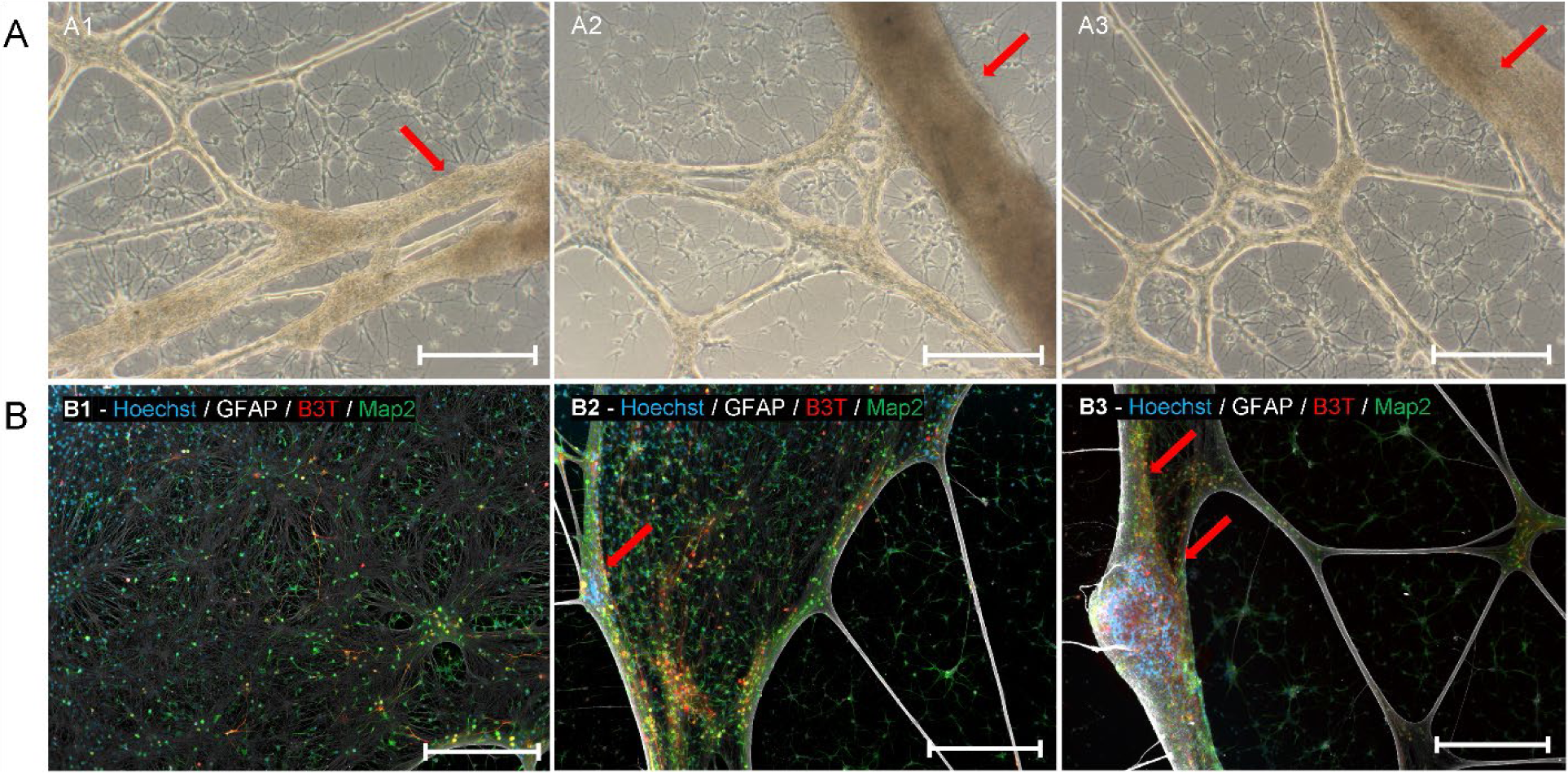
Organization of differentiated uninjured control CC-derived NSCs into large structures. A 1-3: Brightfield image of the formation of larger organized structures (red arrows) after 45 days of differentiation. B1: Unfolded large sheet of neural cells with Beta-3-Tubulin and Map2 positive cells on top of a sheet of GFAP-positive cells. B2: Early stage of folding (red arrow) of neural sheet into large tube-like structures. B3: Late stage of folding and fusing (red arrows) of neural sheet into large tube-like structures. Scale bars A: 50 µm, B: 200 µm.

